# Boosting photosynthetic machinery and defense priming with chitosan application on tomato plants infected with *Fusarium oxysporum* f. sp. *lycopersici*

**DOI:** 10.1101/2020.08.18.256628

**Authors:** Sandra L. Carmona, Andrea del Pilar Villarreal-Navarrete, Diana Burbano-David, Magda Gómez-Marroquín, Esperanza Torres-Rojas, Mauricio Soto-Suárez

## Abstract

Physiological processes of plants infected by vascular pathogens are mainly affected by vascular bundle obstruction, decreasing the absorption of water and nutrients and gas exchange by stomatal closure, and inducing oxidative cascades and PSII alterations. Chitosan, a derivative of chitin present in the cell wall of some organisms including fungi, induces plant defense responses, activating systemic resistance. In this study, the effect of chitosan on the physiological and molecular responses of tomato plants infected with *Fusarium oxysporum* f. sp. *lycopersici* (*Fol*) was studied, evaluating the maximum potential quantum efficiency of PSII photochemistry (Fv/Fm), photochemical efficiency of PSII (Y(II)), photochemical quenching (qP), stomatal conductance (gs), relative water content (RWC), proline content, photosynthetic pigments, dry mass, and differential gene expression (*PAL, LOXA, ERF1,* and *PR1*) of defense markers. A reduction of 70% in the incidence and 91% in the severity of the disease was achieved in plants treated with chitosan, mitigating the damage caused by *Fol* on Fv/Fm, Y(II), and chlorophyll contents by 23%, 36%, and 47%, respectively. Less impact was observed on qP, gs, RWC, and dry mass (16%, 11%, and 26%, respectively). Chitosan-treated and *Fol*-infected plants over-expressed *PR1a* gene suggesting a priming-associated response. These results demonstrate the high potential of chitosan to protect tomato plants against *Fol* by regulating physiological and molecular responses in tomato plants.

## Introduction

*Fusarium oxysporum* is the causative agent of vascular wilt and root rot in plants and is part of a complex of more than 120 *formae speciales* (ff. spp.) that differ in their ability to infect certain hosts (Dean et al., 2012). *F. oxysporum* f. sp. *lycopersici (Fol)* and *F. oxysporum* f. sp*. radicis lycopersici (Forl*) cause significant economic losses in tomato production (Solanki et al., 2015). This pathogen can produce resilient spores called chlamydospores that may persist in the soil for more than 20 years, but are reactivated when the host is detected (Dean et al., 2012; Michielse and Rep, 2009; Di Pietro et al., 2003).

Vascular pathogens such as *Fol* disturb water and nutrient absorption processes in plants, decrease plant growth, and affect transpiration, respiration, and photosynthesis (Yadeta and Thomma, 2013). *Fol* colonizes the elements of the xylem and causes the formation of gums and tylose, obstructing and increasing resistance in rising water, and decreasing the xylem and leaf water potential (Srinivas et al., 2019; Chekali et al., 2011; Lushchak, 2011; Duniway, 1971). Consequently, stomatal closure increases and CO_2_ absorption is reduced in detriment of photosynthetic activity, generating decreases in the quantum efficiency of photosystem II (PSII). These disorders end up affecting the biomass accumulation capacity of the plant and, overall, resemble those caused by water stress (Pinheiro and Chaves, 2011; Chaves et al., 2009; Nogués et al., 2002; Lorenzini et al., 1997).

Light photon reception and electron transport must be regulated to maintain energy production and consumption balance. In high radiation conditions, excess energy must be dissipated to avoid photodamage to the chlorophyll reaction centers (PSI and PSII). Thus, the energy perceived by the PSII can be absorbed and redirected towards photochemical processes such as photolysis and synthesis of adenosine triphosphate (ATP) and nicotinamide adenine dinucleotide phosphate (NADPH^+^); this process is called photochemical quenching (qP). Meanwhile, excess energy is harmlessly dissipated as heat (non-photochemical quenching) or fluorescence, to avoid damage to the leaf. In this way, chlorophyll fluorescence is one of the best indicators for detecting early stress in plants since damage to PSII is avoided, and photosynthetic activity is preserved through energy dissipation as an acclimatization mechanism ( Pérez-Bueno et al., 2019; Azcón-Bieto and Talón, 2008).

The chlorophyll fluorescence parameter most broadly evaluated is the maximum potential quantum efficiency of PSII photochemistry (Fv/Fm); the stability of Fv/Fm value indicates the absence of stress. The photochemical efficiency of PSII (Y(II)) reveals the amount of energy being used in the photochemical phase of photosynthesis, while the photochemical quenching (qP) denotes the ideal (maximum) capacity of the photosystem to receive photons. Thus, a decrease in Y(II) and qP values indicates that the reaction centers are closed (Kardile et al., 2019; Melgarejo et al., 2010; Baker, 2008; Maxwell and Johnson, 2000).

Resistance inducers can be synthetic substances, components derived from plants or microorganisms, as well as Microbial or Pathogen Associated Molecular Patterns (MAMPs or PAMPs) that induce a resistance response in the plant, triggering PTI (PAMP Triggered Immunity) (Ádám et al., 2018; Vidhyasekaran, 2016; Jones and Dangl, 2006). These resistance inducers trigger signaling through hormonal pathways (Salicylic Acid: SA, Jasmonic Acid: JA, and Ethylene: ET), changes in calcium concentrations, ubiquitin-dependent protein degradation, activation of G-proteins, and phosphorylation of Mitogen-Activated Protein Kinase (MAPKs), among others (REF). Then transcription factors or epigenetic modifications regulate the transcriptional activity of pathogenesis-related (PR) genes that encode antimicrobial proteins and antioxidant substances to counteract the oxidative cascades triggered during the defense response (Andersen et al., 2018; Walters et al., 2013; Pitzschke et al., 2009; Shah, 2009).

One of the most studied resistance inducers in plants is chitosan (Orzali et al., 2017), a derivative of chitin present in the cell wall of some fungi, yeasts, green algae, insects, and crustaceans. Chitosan is a linear polymer partially comprising deacetylated N-acetyl glucosamine (GlcNAc) subunits (Orzali et al., 2017; Goy et al., 2009). If more than 50% of the GlcNAc residues are deacetylated at position 2 the heteropolymer is referred to as chitosan (Kappel et al., 2020).

Two mechanisms have been documented by which chitosan protects plants against different phytopathogenic fungi (Lopez-Moya et al., 2019). The first one is the direct action by inhibiting mycelial growth, sporulation, and germination of conidia, mainly through membrane destabilization and cell wall weakening by interrupting the β-1,3 glucan synthesis (Divya et al., 2018; Kumaraswamy et al., 2018; Xing et al., 2018; Zou et al., 2016). The second mechanism is the induction of resistance in plants as a consequence of biochemical and physiological changes such as the production of reactive oxygen species (ROS), the formation of callose deposits, and tissue strengthening (lignification and suberization) (Vidhyasekaran, 2016; Xing et al., 2015; El Hadrami et al., 2010).

Chitosan is widely used worldwide in agriculture through commercial formulations (Table 1), in an international effort to take advantage of a sub-product from the seafood industry, which has great potential to be included as a component of integrated crop management focusing on the minimal use of chemical pesticides and boosting the innate immune system of plants (Ávila-Orozco et al., 2017; Wiesel et al., 2014; Pastor et al., 2013; Pieterse et al., 2013; van Wees et al., 2008; Walters et al., 2005). This work aimed to evaluate the effect of chitosan treatment on tomato plants at the physiological and molecular parameters in during their infection with *Fol*.

**Table 1.**
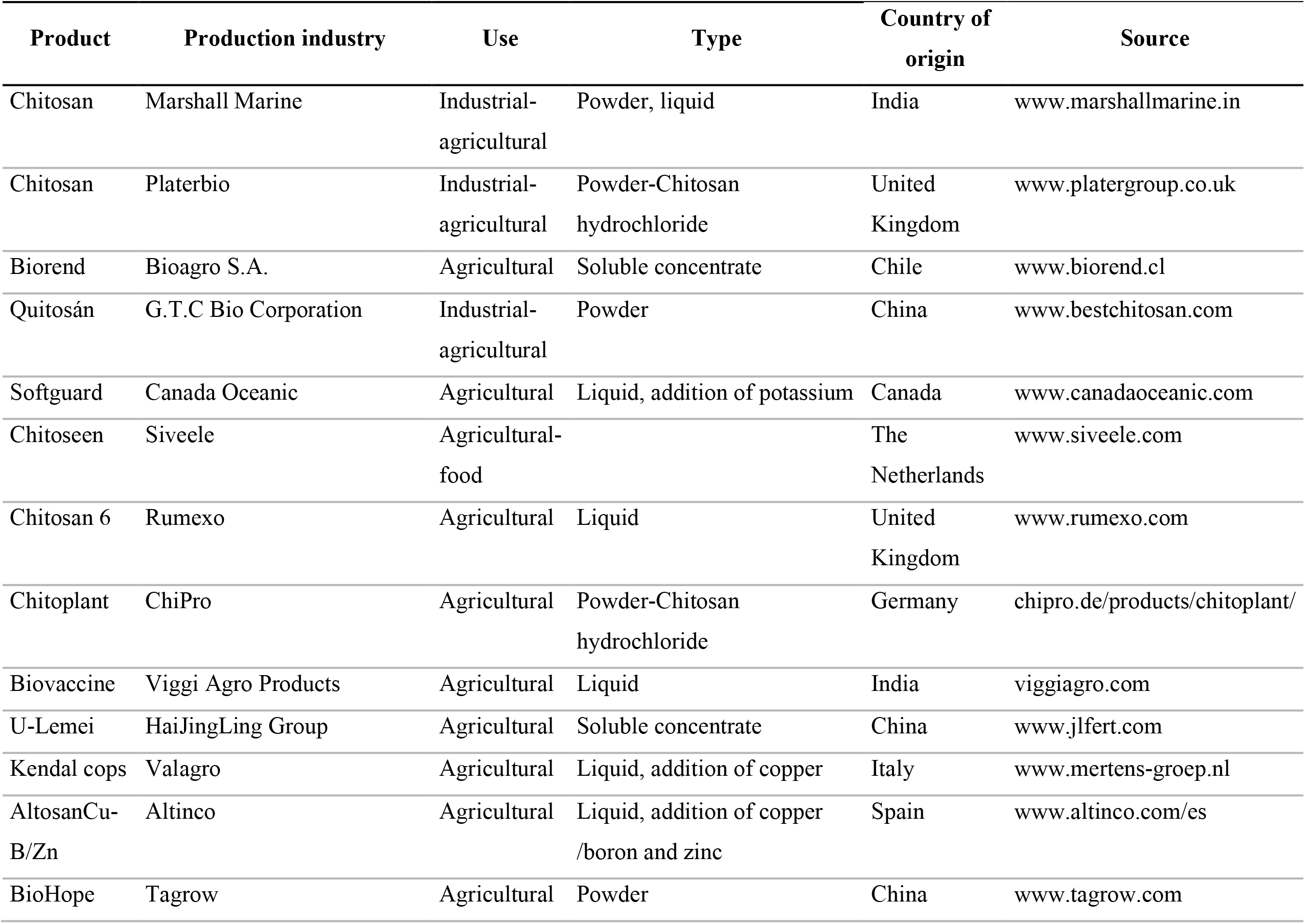
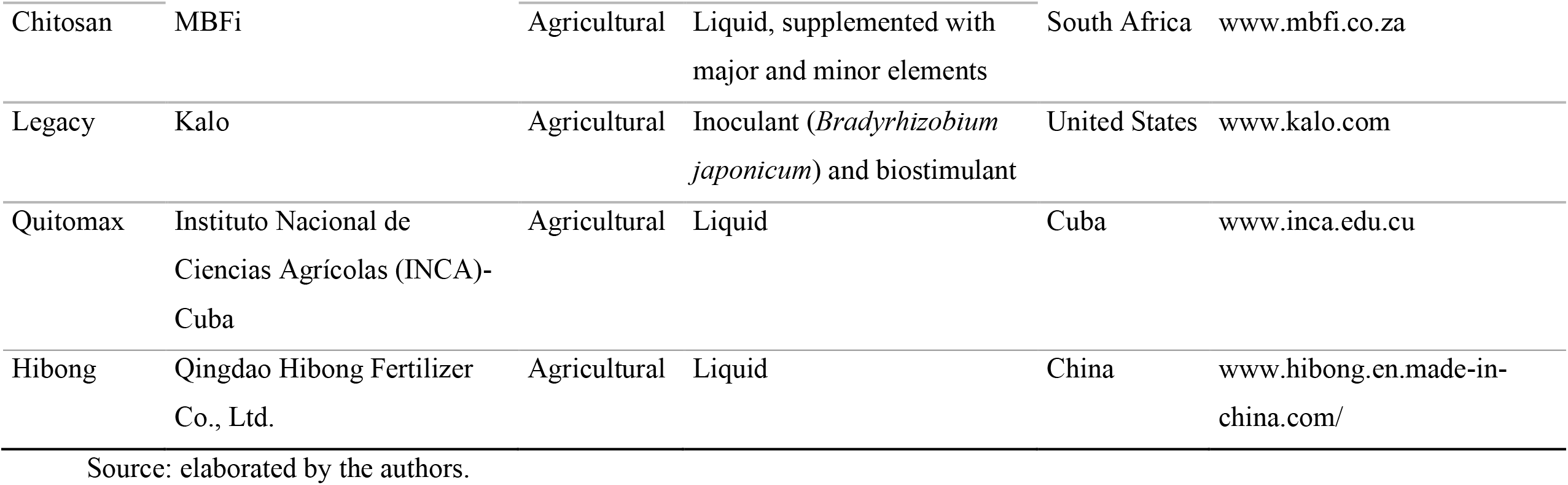
Commercial chitosan-based products available worldwide

## Materials and methods

### Biological material

The *Fol59* strain isolated from tomato plants and identified as *F. oxysporum* f. sp. *lycopersici* (*Fol*) Race 2 (Carmona et al., 2020) was used in all the assays. *Fol59* was mantained on PDA (Potato Dextrose Agar) culture medium added with fresh tomato tissue extract (6 g.L^−1^).

Chonto type tomato seeds of the Santa Cruz Kada variety (Impulsemillas®) were used, superficially disinfected with 2% sodium hypochlorite for 10 min and then with 70% ethanol for one minute. Subsequently, three successive washes with sterile distilled water were carried out, and seeds were germinated on sterile peat. The seedbeds were maintained for 30 days as done on a commercial scale.

### Chitosan preparation

A stock solution of 10 mg.mL^−1^ of chitosan was prepared, adding 2 g of chitosan (Sigma-Aldrich®) in 200 mL of acidified water (1% acetic acid), and the pH was adjusted to 5.6 (Hernández-Lauzardo et al., 2008). From the sterile stock solution elaborated, working solutions were prepared at the concentrations required for each assay.

### *In vitro* inhibition of *Fol59* growth by chitosan

Three molecules of chitosan (Sigma - Aldrich*®*) of different molecular weight (Supplementary Table 1) were used. PDA culture medium was prepared and supplemented with seven concentrations of chitosan (0.5, 1, 1.5, 2, 2.5, 3, and 4 mg.mL^−1^) and a commercial product that was used according to the manufacturer’s recommendations. Unsupplemented PDA (pH 5.6) was used as a negative control.

A 5 mm disk was removed from the margin of14-day-old *Fol59* culture and transferred in the center of a Petri dish with a culture medium supplemented with each chitosan treatment. Five technical replicates were done, and the whole experiment was repeated three independent times (biological replicates). After incubation for seven days at 25 °C, the formula of (You et al., 2016) was used to calculate the IPRG (inhibition percentage (%) on radial growth).

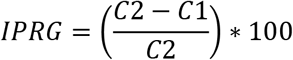

*C2* is the mycelial growth area of *Fol* in the control treatment, and *C1* is the mycelial growth area with chitosan treatment.

### Effect of chitosan on tomato vascular wilt

Thirty-day-old tomato plants were treated with 10 mL of chitosan applied to the soil at different concentrations of each molecular weight considered, 24 hours before transplantation. During the transplant process, plants were inoculated through the root immersion method (Jelinski et al., 2017), using a suspension of *Fol59* of 1,10^6^ con.mL^−1^. Plants immersed in sterile water were used as absolute control, and *Fol59*-infected plants untreated with chitosan were used as pathogenic control.

After transplanting, plants were kept at 30 °C/day and 20 °C/night, with 54% relative humidity under 12 h photoperiod with a light intensity of 90 μmol.m^2.^s^−1^. The variables incidence, severity, and area under the disease progress curve (AUDPC) were evaluated from the onset of symptoms to 14 days after inoculation (DAI). The severity of the disease was established using a modified visual scale (Rongai et al., 2017) from 0 to 5, where 0 is a healthy plant, and 5 is a dead plant. AUDPC was calculated using the following equation:

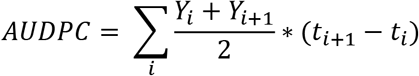

*Y* corresponds to the percentage of the disease, whether expressed as incidence or severity, while *t* is the time elapsed in days (Pedroza and Samaniego, 2009).

### Physiological changes in tomato plants during the interaction with *Fol* and chitosan

For this test, the following treatments were evaluated: (i) Absolute control (plants treated with water), (ii) Chitosan (not infected with *Fol59*), (iii) Chtsn + *Fol* (application of chitosan, and after 24 h infection with *Fol59*), and (iv) Pathogen (Infection with *Fol59*). The concentrations of chitosan used in this experiment were selected according to the *in vitro* and *in planta* assays. Chitosan applications were made 24 h before inoculating the plants with *Fol59*.

### Photochemical efficiency, stomatal conductance, and relative water content

The parameters Fv/Fm, Y(II), and qP were evaluated. In the first case, a miniPAM II modulated fluorometer (Walz Germany®) was used. Stomatal conductance was measured employing an SC-1 porometer (Decagon®, USA). Relative water content (RWC) was calculated using the equation by (Melgarejo et al., 2010) and based on the samples taken at 3, 6, 9, 12, and 15 DAI.

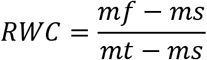

*mf* corresponds to fresh mass, *ms* is dry mass, and *mt* is full turgor mass.

### Proline

Proline content was established using the protocol described by (Ábrahám et al., 2010), taken from Bates et al, (1973), and the absorbance reading was performed at 520 nm employing a microplate spectrophotometer (Biotek®, USES). Proline was calculated using the equation given after calibrating a standard curve: y = 0.0097x + 0.0398, with an R^2^ = 0.98; the results were expressed in μg.g^−1^ of fresh weight.

### Photosynthetic pigments

Carotenoid and chlorophyll a and b contents were calculated in μg.mL^−1^ after extraction using the (Wellburn, 1994) protocol modified by (Rojas-Tapias et al., 2012). Absorbance was detected in a microplate spectrophotometer (Biotek®, USA) using wavelengths of 649 nm and 665 nm for chlorophylls, and 480 nm for carotenoids. The concentration was calculated according to the equations indicated by Wellburn, (1994) as follows:

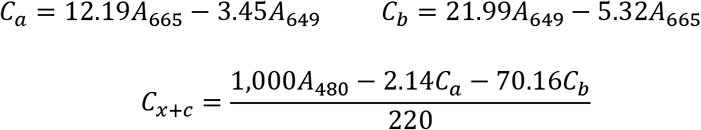

*C*_*a*_ it is chlorophyll a, *C*_*b*_ is chlorophyll b, and *C*_*x*+*c*_ corresponds to carotenoids.

### Dry mass

Complete plants of 9 DAI were dried in an oven at 60 °C for 48 h to establish the accumulation of dry mass.

### Effect of chitosan on the expression of defense marker genes in tomato plants infected with *Fol*

Plants were first treated with chitosan, and 24 h after this treatment, the same plants were infected with *Fol59*, then after 24 h of *Fol59* inoculation (forty-eight hours after chitosan treatment), the foliar part of the plants was collected and flash frozen in liquied nitrogen. Subsequently, total RNA was extracted using the protocol by (Yockteng et al., 2013). The following treatments were assessed: (i) Control (plants treated with water), (ii) Chitosan (not infected with *Fol*), (iii) Chtsn+*Fol* (application of chitosan and subsequent infection with *Fol59*), and (iv) Pathogen (Infection with *Fol59*).

### Assessment of defense and signaling pathway genes

The iScript™ cDNA Synthesis Kit (Bio-Rad®) was used for cDNA synthesis, and the quantitative real-time polymerase chain reaction (qRT-PCR) was performed using the iScript™ One-Step RT-PCR Kit (Bio-Rad®), marking with SYBR® Green (Bio-Rad®), following the manufacturers’ recommendations. Defense-related genes involved in salicylic acid (SA) signaling (*PAL-PHENYLALANINE AMMONIA LYASE* and *PR1a-PATHOGEN RESPONSE1*), and jasmonic acid (JA) signaling (*LOXA-LIPOXYGENASE A*), and the ethylene-response gene (ET) (*ERF1-ETHYLENE RESPONSE FACTOR1*) were evaluated. The tomato gene *EF1a* (elongation factor) was used to normalize the expression. Each reaction was performed in triplicate with two biological replicates. ΔCT values were compared, as described by (Soto-Suárez et al., 2017). The sequence for all primers used in this work are found in Supplementary Table 2.

### Experimental design and data analysis

All experiments were performed separately at least twice, according to a completely randomized block design with a one-plant experimental unit. Measurements were taken every three days until 14 DAI for AUDPC analysis and 15 DAI for physiological parameters. The non-destructive samplings had ten replicates, and the destructive samplings included five replicates on each sampling day. The Statistix 8.0 software was used for data analysis. The normality of the data distribution was verified, and subsequently, an ANOVA and a comparison employing Tukey’s means or the Kruskal-Wallis non-parametric test with α = 0.05 were performed.

## Results

### Effect of chitosan on *Fol* mycelial growth

All the molecules evaluated reduced the radial growth of *Fol59* at a concentration of 1.5 mg.mL^−1^ (Figure 1). As the concentration of the chitosan in the culture medium increased, mycelial growth inhibition was higher (Table 2). Of the three molecules (LMW: low molecular weight, MMW: medium molecular weight, and HMW: high molecular weight), the LMW chitosan inhibited the growth of *Fol59* at the lowest concentrations (0.5 and 1mg.mL^−1^).

**Figure 1.**
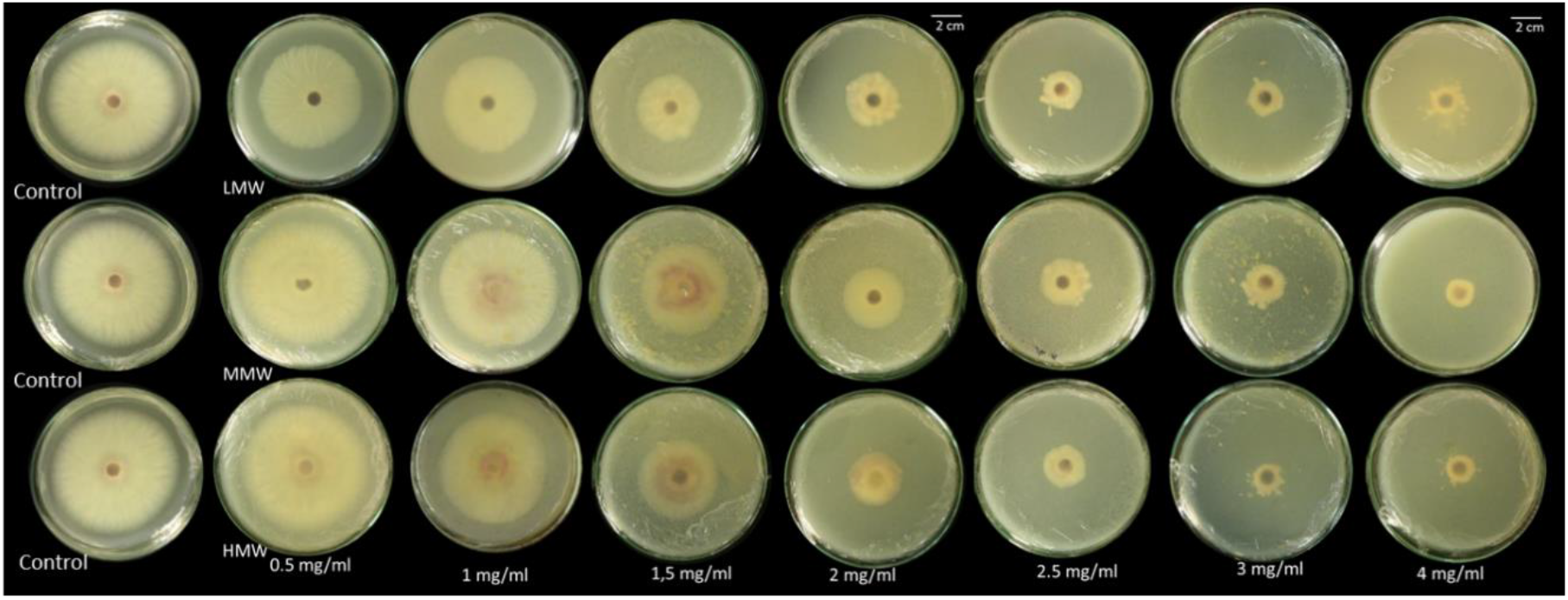
Different chitosan molecules in increasing concentrations inhibit *Fol59* mycelial growth under *in vitro* conditions at seven days after incubation at 25 °C. Source: Photographs taken by the authors.

**Table 2.**
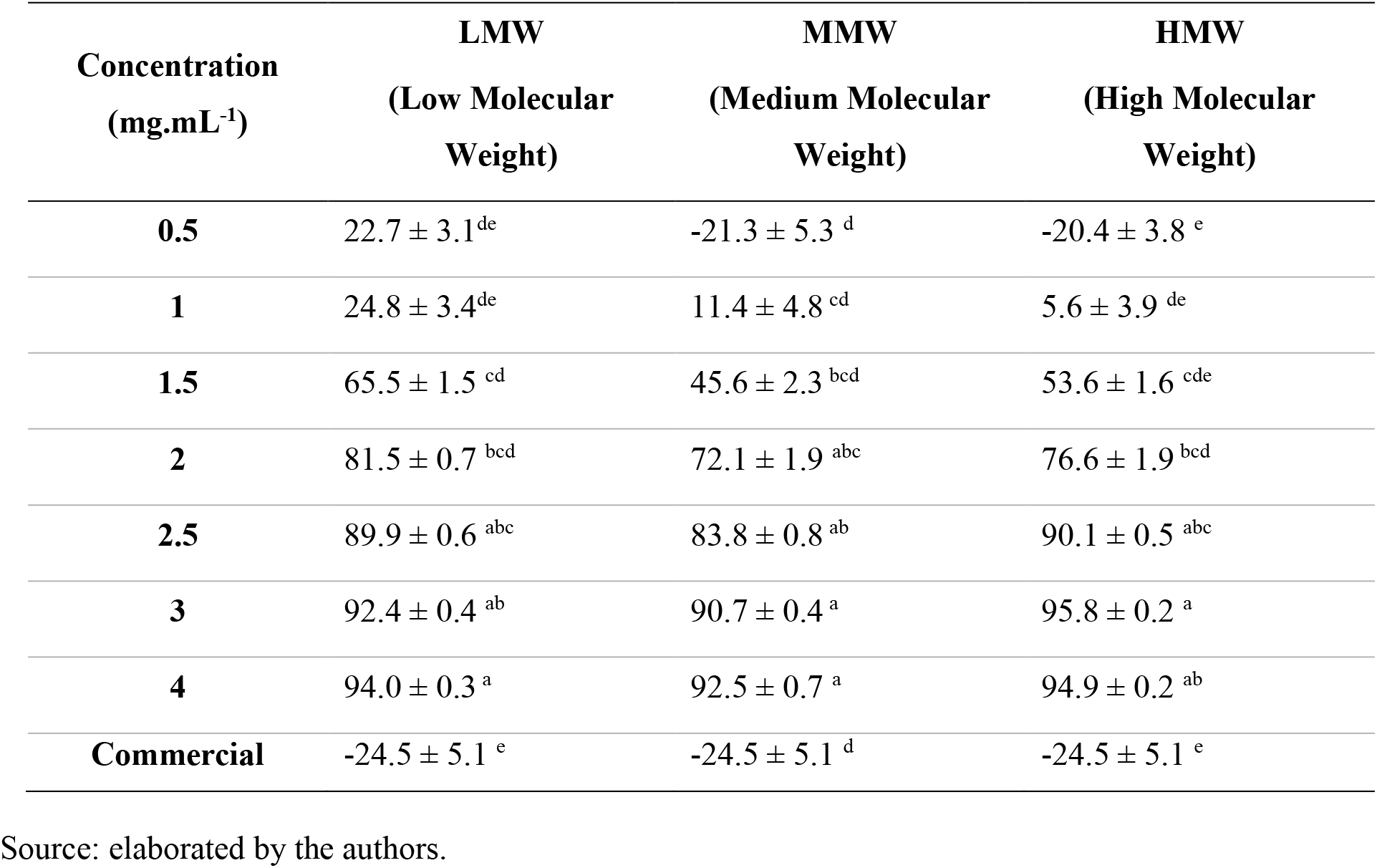
Inhibition percentage on radial growth (IPRG) of *Fol59* after seven days of chitosan application at different concentrations and molecular weights (p = 0.0000; df = 59; F = 66.4). The data are the means of 15 replicates (n = 15). ± corresponds to standard error. Equal letters indicate that there are no statistically significant differences (α = 0.05).

IPRG values in LMW chitosan stabilized above 90% from the concentration of 2.5 mg.mL^−1^. Furthermore, the results obtained for all the molecular weights suggest a correlation between the concentration of chitosan and its antimicrobial capacity (Supplementary Figure 1).

### Effects of chitosan on the reduction of vascular wilt caused by *Fol* in tomato plants

The incidence and severity of the disease in plants treated with LMW and MMW chitosan were significantly lower with the 2.5 and 3 mg.mL^−1^ treatments (Figure 2) in comparison to the other molecules and concentrations. However, the best performance in terms of incidence efficacy was obtained by LMW chitosan in concentrations of 2.5 and 3 mg.mL^−1^ (70% and 67.5%, respectively). Regarding the efficacy percentage on severity, the best treatment reached 91% (LMW: 2.5 mg.mL^−1^), followed by 85% (LMW: 3 mg.mL^−1^) and 78% (MMW: 3 mg.mL^−1^) (Table 3). In agreement with what was found *in vitro*, the low concentrations of chitosan were not able to effectively decrease the expression of the disease.

**Figure 2.**
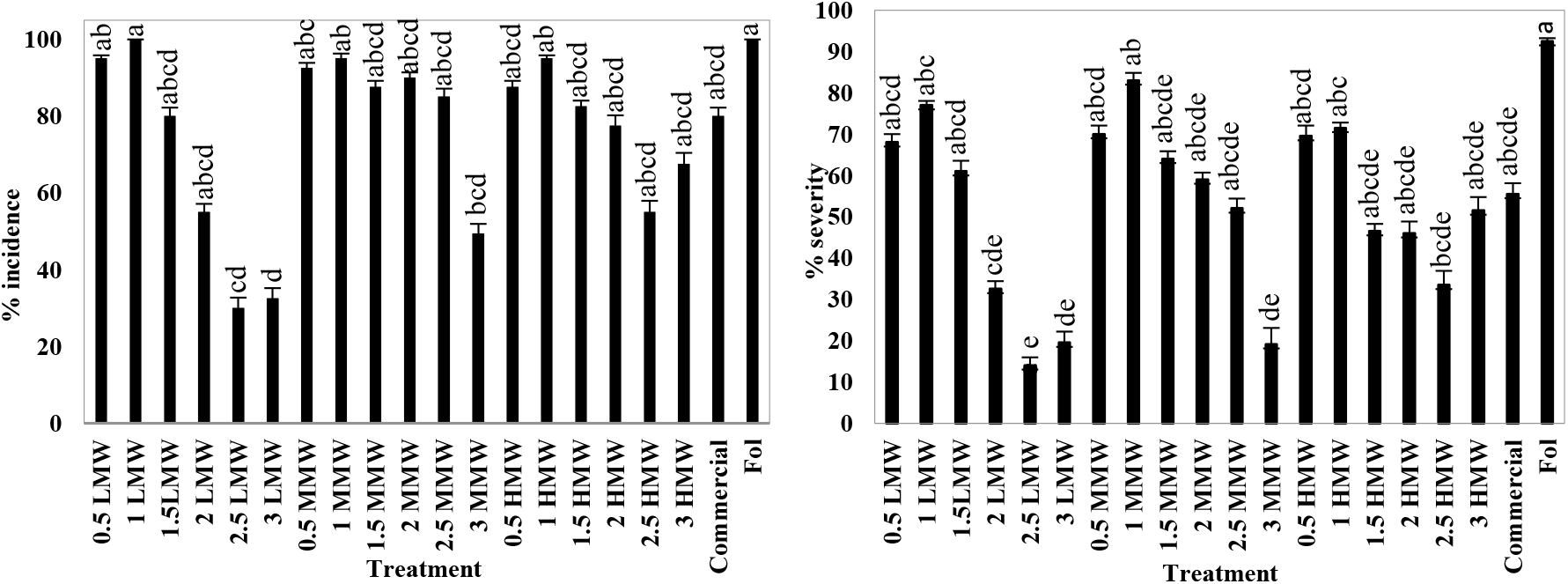
Results of the effect of chitosan on the incidence (α = 0.05; p = 0.0000; F = 8.68; df = 159) and severity (α = 0.05; p = 0.0000; F = 10.2; df = 159) of the disease 14 DAI in terms of percentage. The data are means of 40 replicates (n = 40). The bars correspond to the standard error. Equal letters indicate that there are no statistically significant differences (α = 0.05). Source: elaborated by the authors.

**Table 3.** Effect of chitosan on the AUDPC incidence (p=0.000; F=13.4; df=159) and severity (p=0.000; F=16.5; df=159), and the efficacy on incidence (p=0.000; F=8.20; df=159) and severity (p=0.000; F=11.4; df=159) in infected plants with *Fol59.*

| Molecule | Concentration<br>(mg.mL<sup>-1</sup>) | Incidence |  | Severity |  |
| --- | --- | --- | --- | --- | --- |
|  |  | AUDPC | %<br>effectiveness | AUDPC | %<br>effectiveness |
| LMW (Low<br>Molecular<br>Weight) | 0.5 | 390±4.1<sup>abcdef</sup> | 5.0±3.9<sup>bc</sup> | 196±3.5<sup>abcdef</sup> | 20.6±3.3<sup>bcd</sup> |
|  | 1 | 670±8.8<sup>abc</sup> | 0.0±0.0<sup>c</sup> | 318±4.3<sup>abc</sup> | 0.1±9.0<sup>cd</sup> |
|  | 1.5 | 360±5.2<sup>abcdef</sup> | 20.0±5.0<sup>abc</sup> | 181±3.9<sup>abcdef</sup> | 33.3±3.6<sup>abcd</sup> |
|  | 2 | 210±5.2<sup>def</sup> | 45.0±4.3<sup>abc</sup> | 84±3.8<sup>def</sup> | 61.2±2.8<sup>abc</sup> |
|  | 2.5 | 80±6.0<sup>f</sup> | 70.0±5.5<sup>a</sup> | 28±3.8<sup>f</sup> | 90.9±1.2<sup>a</sup> |
|  | 3 | 113±13.7<sup>ef</sup> | 67.5±2.2<sup>a</sup> | 45±7.1<sup>ef</sup> | 85.0±1.3<sup>ab</sup> |
| MMW<br>(Medium<br>Molecular<br>Weight) | 0.5 | 583±7.1<sup>abc</sup> | 7.5±5.1<sup>bc</sup> | 284±5.7<sup>abcd</sup> | 7.9±3.0<sup>cd</sup> |
|  | 1 | 715±3.0<sup>ab</sup> | 5.0±5.9<sup>bc</sup> | 371±2.9<sup>ab</sup> | -17.0±8.2<sup>d</sup> |
|  | 1.5 | 498±15.7<sup>abcdef</sup> | 12.5±4.8<sup>abc</sup> | 223±11.1<sup>abcdef</sup> | 14.6±6.1<sup>abcd</sup> |
|  | 2 | 530±3.9<sup>abcd</sup> | 10.0±4.5<sup>abc</sup> | 237±3.3<sup>abcde</sup> | 26.2±2.4<sup>abcd</sup> |
|  | 2.5 | 341±9.2<sup>bcdef</sup> | 15.0±5.6<sup>abc</sup> | 155±6.3<sup>bcdef</sup> | 43.0±3.7<sup>abcd</sup> |
|  | 3 | 291±17.9<sup>bcdef</sup> | 50.6±2.1<sup>ab</sup> | 79±8.1<sup>def</sup> | 78.4±1.7<sup>ab</sup> |
| HMW (High<br>Molecular<br>Weight) | 0.5 | 503±6.1<sup>abcde</sup> | 12.5±4.8<sup>abc</sup> | 296±6.5<sup>abcd</sup> | 7.4±5.3<sup>cd</sup> |
|  | 1 | 695±7.0<sup>abc</sup> | 5.0±3.9<sup>bc</sup> | 353±4.4<sup>abc</sup> | -2.8±7.7<sup>cd</sup> |
|  | 1.5 | 378±8.8<sup>abcdef</sup> | 17.5±3.7<sup>abc</sup> | 155±5.7<sup>bcdef</sup> | 51.5±3.0<sup>abcd</sup> |
|  | 2 | 303±11.0<sup>bcdef</sup> | 22.5±5.7<sup>abc</sup> | 132±9.2<sup>bcdef</sup> | 58.6±2.8<sup>abcd</sup> |
|  | 2.5 | 265±6.6<sup>cdef</sup> | 45.0±4.4<sup>abc</sup> | 120±5.1<sup>cdef</sup> | 60.3±3.8<sup>acd</sup> |
|  | 3 | 363±6.1<sup>abcdef</sup> | 32.5±5.9<sup>abc</sup> | 180±6.1<sup>abcdef</sup> | 55.9±4.1<sup>abcd</sup> |
|  | Commercial | 350±7.3<sup>bcdef</sup> | 20.0±4.5<sup>abc</sup> | 164±5.8<sup>abcdef</sup> | 37.0±2.4<sup>abcd</sup> |
|  | Pathogen | 915±4.3<sup>a</sup> | 0±0.0<sup>bc</sup> | 504±3.8<sup>a</sup> | 0±0.0<sup>cd</sup> |
|  | Control | 0±0.0<sup>f</sup> | 100.0±0.0<sup>a</sup> | 0±0<sup>f</sup> | 100.0±0<sup>a</sup> |

The data are means of 40 replicates (n = 40). ± corresponds to standard error. Equal letters indicate that there are no statistically significant differences ( $\alpha = 0.05$ ). Source: elaborated by the authors.

Chitosan also significantly reduced disease progression over time, reflected in the AUDPC variable (Table 3), demonstrating its ability to delay the effects of the disease compared to infected untreated plants. The results showed the protective effect of chitosan against vascular wilt (Figure 3), and the best performance of the LMW (2.5 mg.mL^−1^) treatment; therefore, it was selected for the following tests.

**Figure 3.**
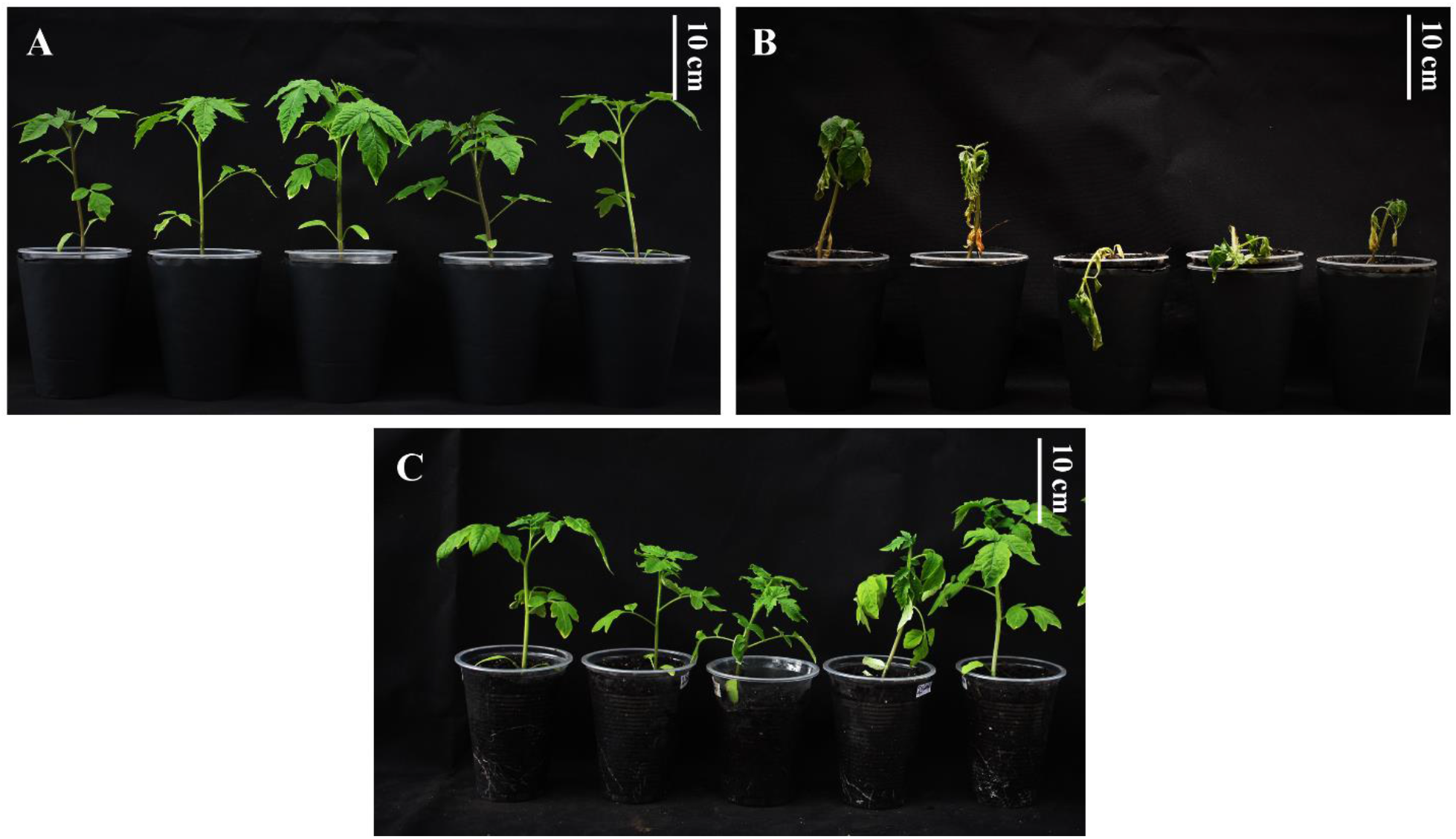
Visual symptoms of vascular wilt 14 DAI. A) Absolute control; B) Pathogen control (infected with *Fol*); C) Plants treated with chitosan (Low Molecular Weight: 2.5 mg.mL^−1^) and inoculated with *Fol59*. Source: elaborated by the authors.

### Effect of chitosan on physiological parameters of tomato plants infected with *Fol*

The results on Fv/Fm, Y(II) and qP, indicate a reduction in photosynthetic processes due to the infection by *Fol*59 (Figure 4, 5, and 6). To investigate if chitosan can mitigate the early tomato stress conditions caused by *Fol*59, we measured photosynthesis effiency (Fv/Fm). In this experiment, the Fv/Fm decreased in the infected plants from 9 DAI, observing a decrease of up to 70% at 15 DAI in plants inoculated with *Fol*59 compared to uninfected control plants (Figure 4). In contrast, in the Chtsn + *Fol* treatment (plants applied with chitosan 24 hours before being infected with *Fol*), the Fv/Fm was 7.5% lower than the absolute control (control). The plants infected only with *Fol* (pathogenic treatment), the Fv/FM were significantly different (p<0.05).

**Figure 4.**
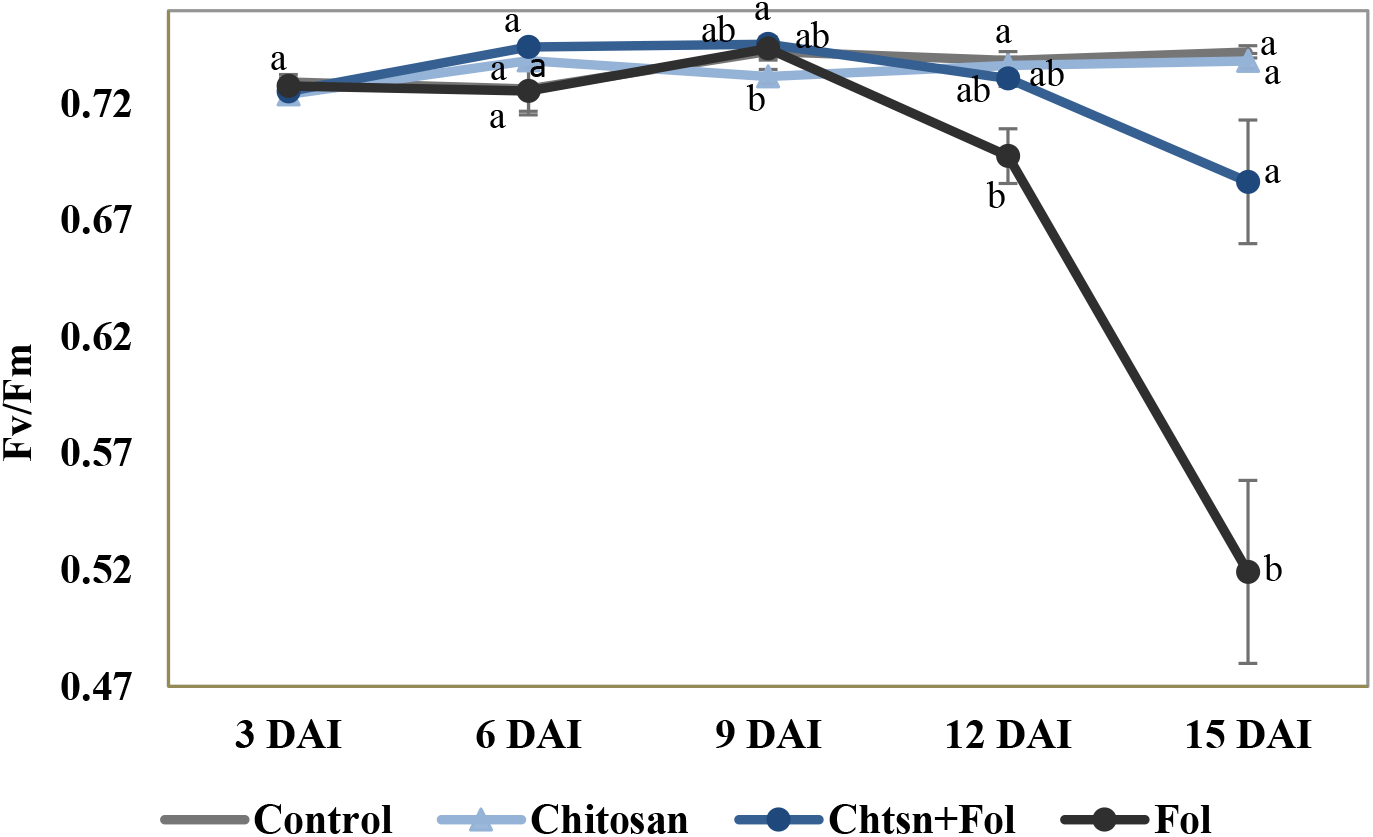
Fv/Fm progress during 15 days in tomato plants with different treatments (p = 0.000; F = 15.3; df = 79). Control (plants treated with water); Chitosan (not infected with *Fol*), Chtsn + *Fol* (application of chitosan and subsequent infection with *Fol*); *Fol* (Infected with *Fol59*). The data are means of 20 replicates (n = 20). The bars correspond to the standard error. Equal letters indicate that there are no statistically significant differences (α = 0.05). Source: elaborated by the authors.

**Figure 5.**
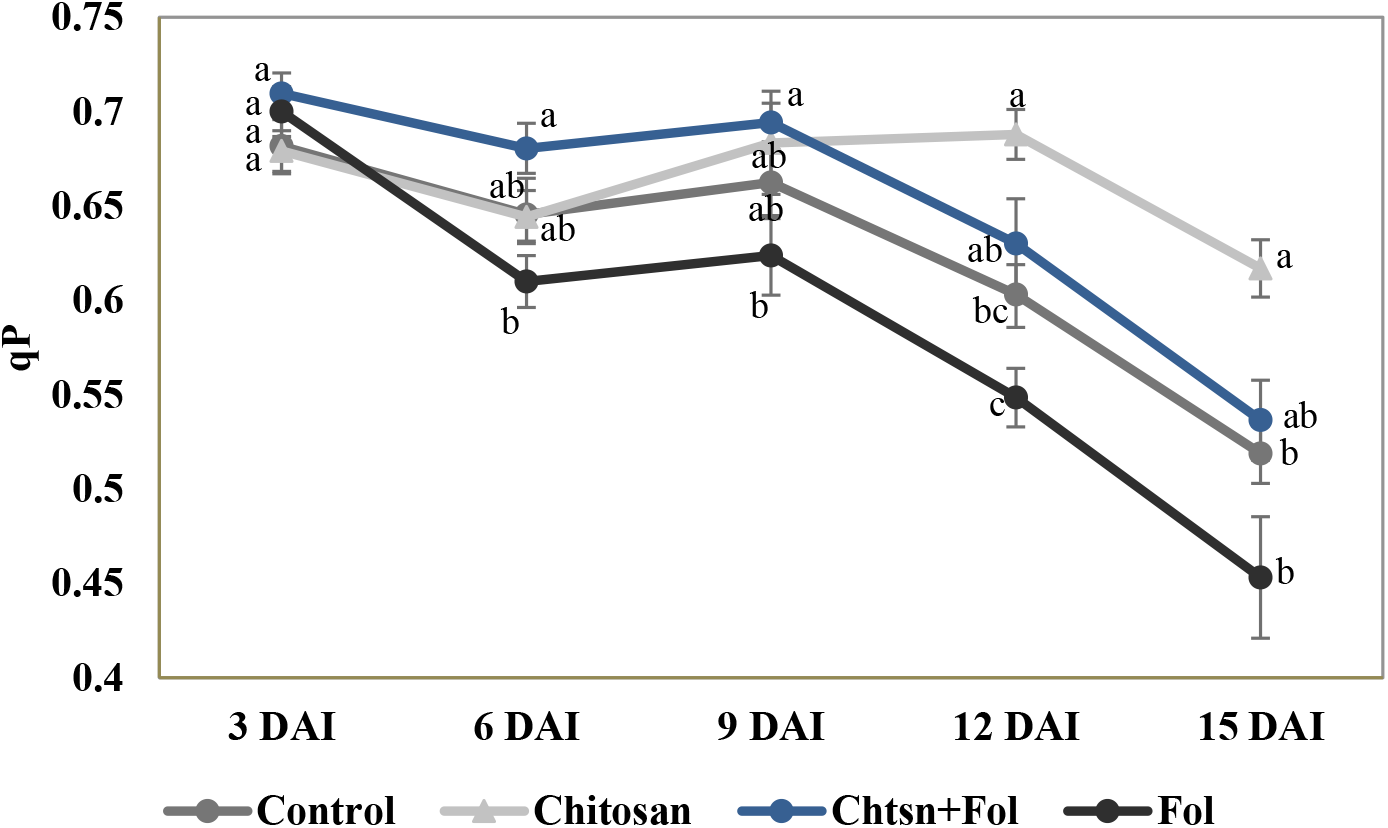
qP progress during 15 days in tomato plants with different treatments (p = 0.0001; F = 8.55; df = 79). Control (plants treated with water); Chitosan (not infected with *Fol59*), Chtsn + *Fol* (application of chitosan and subsequent infection with *Fol59*); *Fol* (Infected with *Fol59*). The data are the means of 20 replicates (n = 20). The bars correspond to the standard error. Equal letters indicate that there are no statistically significant differences (α = 0.05). Source: elaborated by the authors.

**Figure 6.**
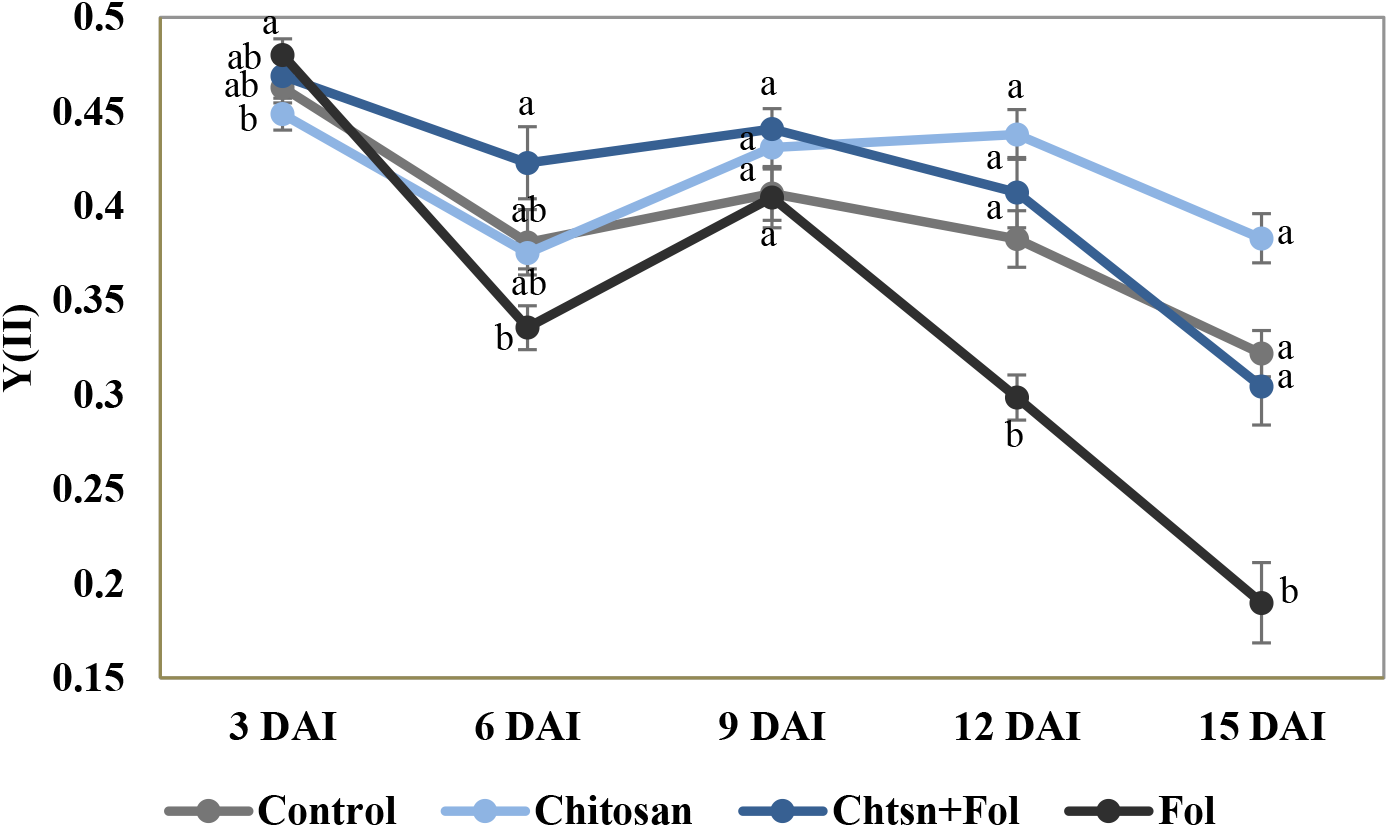
Y(II) progress during 15 days in tomato plants with different treatments (p = 0.000; F = 18; df = 79). Control (plants treated with water); Chitosan (not infected with *Fol59*), Chtsn + *Fol* (application of chitosan and subsequent infection with *Fol59*); *Fol* (Infected with *Fol59*). The data are the means of 20 replicates (n = 20). The bars correspond to the standard error. Equal letters indicate that there are no statistically significant differences (α = 0.05). Source: elaborated by the authors.

As qP indicates the photochemical photosynthesis phase, it was used to investigate the photosynthetic activity induced by *Fol59* and chitosan application. According to our results, the qP was highly sensitive since it decreased early (6 DAI) in the *Fol59* treatment (Figure 5). Plants infected with *Fol* showed a 15.5% decrease in qP (0.45) compared to plants previously treated with chitosan and then infected with *Fol* (Chtsn + *Fol59*), the latter showing a qP value of 0.51. However, the qP values for both treatments did not show significant differences.

Parameter Y(II) responded similarly to qP, decreasing at 6 DAI in the *Fol* treatment, although with a slight rebound on day nine after inoculation (Figure 6). Subsequently, the Y(II) value for the plants infected with *Fol59* decreased significantly. Lower Y(II) values represent decreases in the energy flow destined for the ATP and NADPH production. Plants previously treated with chitosan and then infected with *Fol59* (Chtsn + *Fol*) showed a lower decrease in Y(II) values (5%) compared to plants only infected with *Fol59* (41%).

The results obtained in Fv/Fm and Y(II) suggest an important role of the treatment with chitosan, mitigating the damage by *Fol* on the light phase of tomato photosynthesis.

### Stomatal conductance

One of the most well-known alterations produced during *Fol* infection is the difficulty by the plant in absorbing water and nutrients (Yadeta and Thomma, 2013), inducing stomatal closure. As a result, in this study, plants infected with *Fol59* showed a 64% decrease in gs than the control plants, indicating an increase in resistance to xylem water flow. At 9 DAI, a 40% decrease in gs in the *Fol* treatment compared to the Chtsn + *Fol* treatment (Supplementary Table 3) is explained by the proliferation of the pathogen in the xylem. This trend was maintained until 15 DAI, where the gs was 87% lower in the *Fol* treatment plants compared to the control treatment; meanwhile, in the Chtsn + *Fol* treatment, the reduction was 57% lower compared to the control (Table 4). Despite not detecting significant differences between the *Fol* and Chtsn + *Fol* treatments, it seems that chitosan-treated plants show less hydric stress and less stomatal closure than the plants only infected with *Fol59*. This results are suggesting that chitosan treatment may reduce water limitations by not closing the same proportion of stomata.

**Table 4.** Response of the variables gs (p = 0.000; F = 35.2; df = 39), RWC (p = 0.0376; F = 3.13; df = 39), photosynthetic pigments (p = 0.0008; F = 6.94; df = 39), and proline content (p = 0.000; F = 10.4; df = 39) to the treatments assessed. Control (plants treated with water); Chitosan (not infected with *Fol59*), Chtsn + *Fol* (application of chitosan and subsequent infection with *Fol59*); *Fol* (infected with *Fol59*).

| Treatment | gs | RWC | Chlorophyll a | Chlorophyll b | Carotenoids | Proline |
| --- | --- | --- | --- | --- | --- | --- |
| **Control** | 272.5±30.97<sup>a</sup> | 0.746±0.05<sup>ab</sup> | 6±0.15<sup>a</sup> | 2.1±0.05<sup>a</sup> | 0.66±0.02<sup>a</sup> | 6.1±0.69<sup>b</sup> |
| **Chitosan** | 290.8±28.00<sup>a</sup> | 0.810±0.04<sup>a</sup> | 6.8±0.14<sup>a</sup> | 2.4±0.04<sup>a</sup> | 0.82±0.03<sup>a</sup> | 4.8±0.72<sup>b</sup> |
| **Chitosan+*Fol*** | 131.9±20.54<sup>ab</sup> | 0.671±0.05<sup>ab</sup> | 6.5±0.15<sup>a</sup> | 2.2±0.05<sup>a</sup> | 0.88±0.02<sup>a</sup> | 38.6±8.35<sup>a</sup> |
| **Pathogen** | 33±3.56<sup>b</sup> | 0.592±0.05<sup>b</sup> | 3.6±0.19<sup>b</sup> | 1.3±0.06<sup>b</sup> | 0.59±0.03<sup>a</sup> | 54.4±9.02<sup>a</sup> |

The data are means of 10 replicates (n = 10). ± corresponds to standard error. Equal letters indicate that there are no statistically significant differences ( $\alpha = 0.05$ ). Source: elaborated by the authors.

### Relative water content (RWC)

The RWC remained stable in the first phase of the infection (6 DAI) and began to decrease at 9 DAI in plants infected with *Fol59* (Supplementary Table 3). It is important to note that after 9 DAI, the RWC was 10% higher in the chitosan treatment compared to the absolute control, suggesting an optimization of the water use may be induced by chitosan. However, there were no significant differences between the inoculated treatments; therefore, in the Chtsn + *Fol* treatment plants, this effect does not seem sufficient to mitigate the water limitations caused by the pathogen.

### Proline content

At 15 DAI, the proline content in the plants infected with *Fol59* was nine times higher than the control, and in the Chtsn + *Fol* treatment, the values were six times higher compared to the control. The proline values were significantly higher in the treatments inoculated with *Fol59* compared to those not inoculated (Table 4).

### Chlorophyll content

The chlorophyll content was stable until 12 DAI. From 15 DAI in the plants of the pathogenic treatment, chlorophylls a and b showed a decrease of 38% to the control; meanwhile, in the Chtsn + *Fol* treatment, the contents remained stable without significant differences compared to the control (Table 4). The carotenoid content was not altered in any of the treatments during the experiment. The decrease in chlorophyll concentrations is consistent with the appearance of chlorosis in plants only infected with *Fol*.

### Dry mass

The accumulation of biomass in the plant under stress conditions depends on the adequate use of energy for direct metabolic and photosynthetic processes, constituting an indicator of plant health (Figure 7). To gain an insight into the effect of the chitosan application on tomato plant health infected with *Fol59,* the accumulation of biomass was evaluated.

**Figure 7.**
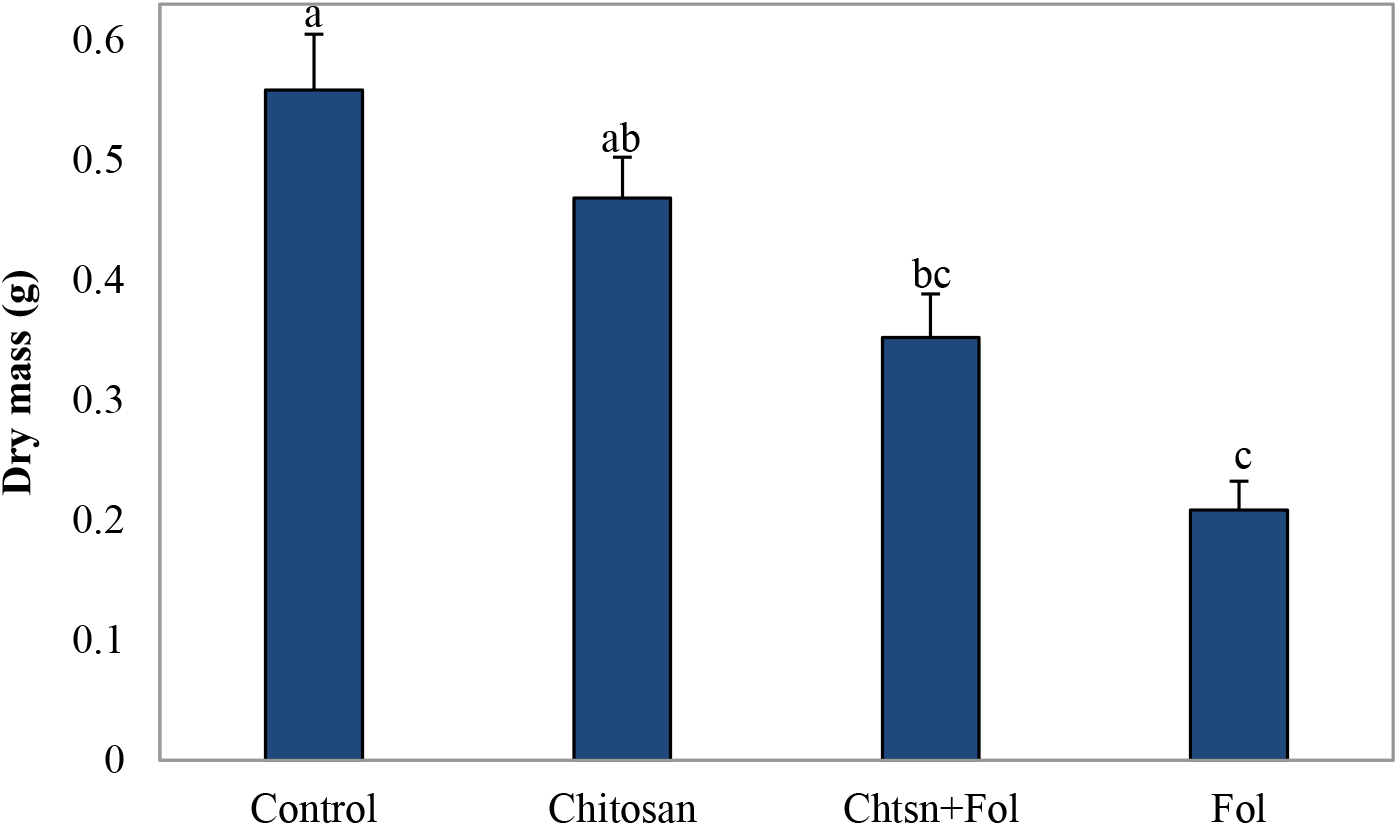
Dry mass accumulation in tomato plants with different treatments (p = 0.000; F = 20; df = 79) 9 DAI. Control (plants treated with water); Chitosan (not infected with *Fol59*), Chtsn + *Fol* (application of chitosan and subsequent infection with *Fol59*); *Fol* (Infected with *Fol59*). The data are the means of 20 replicates (n = 20). The bars correspond to the standard error. Equal letters indicate that there are no statistically significant differences (α = 0.05). Source: elaborated by the authors.

Our results indicate that 9 DAI, *Fol59* infection significantly affects the biomass accumulation in plants, decreasing from 36% and 62% in the Chtsn + *Fol* and *Fol* treatments, respectively, compared to the control. The two treatments with *Fol59* inoculated plants did not differ significantly between them.

### Effect of chitosan on the expression of defense marker genes in tomato plants infected with *Fol*

The differential expression of the *PR1a, ERF1, PAL,* and *LOXA* genes in leaves and stems was evaluated after 72 hours of chitosan application and 48 hours after infection with *Fol59* (Figure 8). In the chitosan treatment (without *Fol59* inoculation), the *ERF1, LOXA,* and *PAL* genes were not differentially expressed compared to the control, while the *PR1a* gene was expressed 3.6 times more compared to the control.

**Figure 8.**
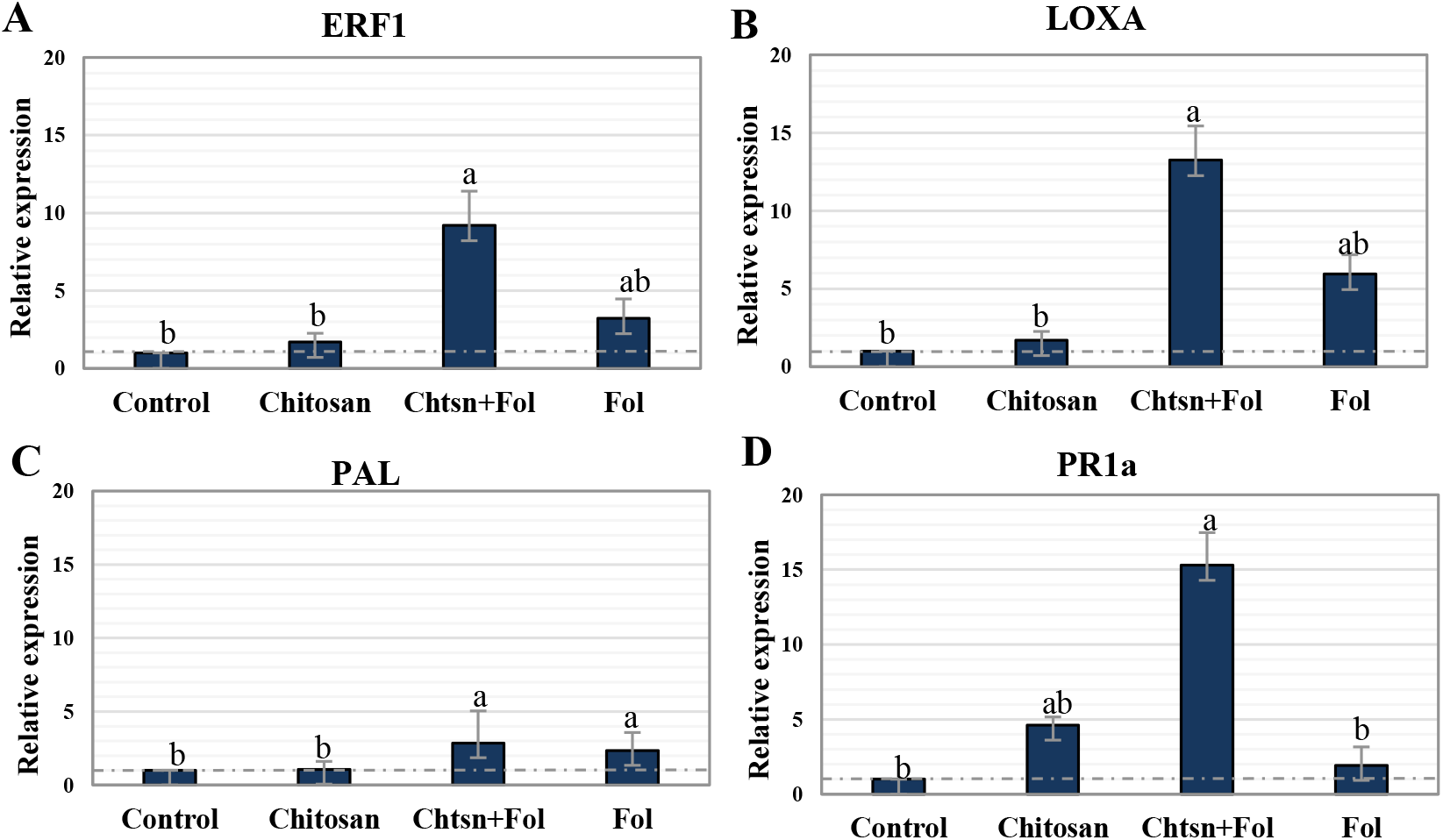
Differential expression of the defense genes A) *ERF1* (ET/JA) (p = 0.0002; F = 16.1; df = 17); B) *LOXA* (JA) (p = 0.0000; F = 30.3; df = 17); C) *PAL* (SA) (p = 0.0002; F = 16.1; df = 17), and D) *PR1a* (SA) (p = 0.0221; F = 4.96; df = 17) in tomato plants subjected to different treatments. Standardized data with respect to the control (= 1). Control (plants treated with water); Chitosan (not infected with *Fol59*), Chtsn + *Fol* (application of chitosan and subsequent infection with *Fol59*); *Fol* (Infected with *Fol59*). The data are the means of 6 replicates (n = 6). The bars correspond to the standard error. Equal letters indicate that there are no statistically significant differences (α = 0.05). Source: elaborated by the authors.

For the Chtsn + *Fol* treatment, all genes were differentially expressed compared to the control. The *PAL, ERF1*, *LOXA*, and *PR1a* genes were expressed 1.8, 8.2, 12.2, and 14.2 times more, respectively, compared to the control plants. This results suggests that 72 hours after applying chitosan and 48 hours after *Fol59* inoculation, detection of the pathogen activates a systemic response that extends to the aerial part of the plant, even when the infection has not spread to that level. On the other hand, in the *Fol* treatment, the *PR1a* gene did not differ from the control (0.9 times more than the control), while the *ERF1, LOXA,* and *PAL* genes were expressed 2.2, 4.9, and 1.3 times more than the control.

## Discussion

The high *in vitro* growth inhibition of chitosan on *Fol59* is probably due to the antimicrobial mechanisms of chitosan on fungi, affecting mainly cell walls by inhibition in glucan biosynthesis and destabilization of cell membranes (Kumaraswamy et al., 2018). In this sense, other authors confirmed that chitosan produces an increase in potassium ion flow in microbial cells and that the increase in chitosan concentrations proportionally increases the rapid loss of these electrolytes (Xing et al., 2015).

In our research, chitosan exerts antifungal activity *in vitro*. These results coincide with those reported by Oh et al. (2019), who found that both chitosan and nanoparticles derived from it can be used effectively against phytopathogenic fungi when evaluating the antifungal and antibacterial action in tomato phytopathogens. Conversely, Soliman and El-Mohamedy, (2017) and El-Mohamedy et al. (2019) found that HMW and LMW chitosan effectively inhibited the growth of fungal pathogens and that this inhibition increased depending on the concentration. Besides, the results indicated that growth and sporulation decreased significantly when the LMW molecule was used compared to the HMW molecule.

The results of this work are in agreement with several authors such as Sathiyabama and Charles (2015) and Saharan et al. (2015) who applied chitosan on tomato plants, finding reductions of 81% and 61% in vascular wilt, and Lafontaine and Benhamou, (1996) who observed 90% reduction in crown rot due to *Forl*. It demonstrates the protective effect of chitosan on tomato plants against vascular wilt caused by *Fol*. The efficacy of LMW chitosan was consistent *in vitro* and *in planta* assays and, in accordance with other studies for plant pathogens, including *F. oxysporum* (Zhang et al., 2017; Alburquenque et al., 2010; Tikhonov et al., 2006), highlighting the physicochemical properties of chitosan and its ability to induce resistance in plants against phytopathogens (Orzali et al., 2017). The process is presumed to occur by binding the chitosan molecule to the plant cell membrane, initiating signaling by hydrogen peroxide (H_2_O_2_) and nitric oxide (NO) in photosynthetic plastids (Kauss et al., 1989), triggering antioxidant substance synthesis and abscisic acid (ABA), which ends in the induction of stomatal closure, gene activation and other responses related to biotic stress (Zhang, 2004).

Fv/Fm has been widely used as a stress indicator in plants (including tomato), and its values decrease according to the severity of the stress. This parameter represents a measure of the photon absorption capacity by PSII intended to reduce plastoquinone A (Zhou et al., 2015; Kalaji et al., 2012). Moreover, this value is a sensitive indicator of plant photosynthesis, whose optimal values are around 0.83 in stress-free plants (standard for a wide variety of plant species) (Maxwell and Johnson, 2000). Plants inoculated with *F. oxysporum,* with a severity level of 92%, (Wagner et al., 2006) reported a decrease in Fv/Fm to 0.343, highlighting the relationship between this variable and disease stress. In this work, the fact of not finding significant differences in the Fv/Fm in untreated plants (absolute control) compared to plants only treated with chitosan, reinforces the potential of using chitosan as an alternative *Fol* control without the appearance of phytotoxic effects.

Decreases in Fv/Fm values are a clear indicator of severe stress that is causing damage to the photosynthetic apparatus (Goltsev et al., 2016). In this study, the first symptoms of wilt were observed 6 DAI; while alterations in Fv/Fm were detected 12 DAI, simultaneously with the appearance of symptoms on the leaves evaluated (upper third). These results indicate that during early infection, no damage to PSII is detected in the upper leaves. The decrease in Fv/Fm is associated with photochemical damage since the reduction in the number of active reaction centers indicates a lower use of the photons that hit the light-collecting center, caused by the damage of the disease.

On the other hand, lower qP values 6 DAI in plants with *Fol* show that a decrease in the flow of electrons between photosystems occurs, probably due to disorders in membranes and proteins in the chloroplast (ferredoxins and quinones, among others). Low qP values after 15 days in *Fol* inoculated plants suggest extensive damage to the PSII reaction centers caused by *Fol*. In contrast, this impact was slightly lower on the other treatments. This study suggests that qP is a more sensitive variable than Fv/Fm as it decreases earlier during infection. This may be related to the energy dissipation mechanism that is not intended for photosynthesis, which instead of dissipating in the form of fluorescence, may have dissipated in the form of heat.

Nogués et al. (2002) found affectations of 28% and 27% in qP and Fv/Fm, respectively, as well as Segarra et al. (2010), who described decreases of 58% and 24% in qP and Fv’/Fm’, respectively, after 31 days of inoculation with *Fol*. In both cases, the variable qP seems to be more sensitive and indicates a low regulation in electron transport. These authors attribute this to the reduction in the uptake and assimilation of CO_2_ in leaves caused by stomatal closure and increased xylem resistance, hindering water absorption. Furthermore, the authors argue that photoinhibitory damage may be related to low demand for cations and ATP, causing the reaction centers to close.

In this work, the reduction in Fv/Fm occurred after 12 DAI, long after the first symptoms appeared (6 DAI), suggesting that in the initial stages of infection, no irreversible damage occurs in the PSII and that this happens as a consequence of the xylem obstruction by the pathogen and a decrease in stomatal conductance.

Furthermore, according to our results, *Fol* seriously compromises the photochemical processes of photosynthesis through the energy flow. This result is similar to the one reported by Nogués et al. (2002), where *Fol* infection decreased by 50% the Y(II) in tomato plants. Besides, decreases in Fv/Fm and electron transfer rate (ETR, Supplementary Figure 2) were found in the current study, leading to a decrease in ribulose-1,5-bisphosphate carboxylase/oxygenase (Rubisco) activity. In this sense, Pshibytko et al. (2006) attributed this result to a decrease in electron transport due to decreases in the acceptor flux in quinone A (QA) of the PSII.

The symptoms caused by *Fol* infection have previously been related to disorders caused by water stress (Duniway, 1971); for example, decreases in the photosynthetic rate of infected plants have been correlated with the consequences of the decrease in gs, such as the water state and gas exchange (CO_2_ absorption) ( Nogués et al., 2002; Lorenzini et al., 1997). These alterations can generate the accumulation of ROS and a decrease in the rate of CO_2_ assimilation in the chloroplast, and induce photorespiration and damage to thylakoid membranes, affecting the PSII, which partly explains the results in the variables Fv/Fm and Y(II) in the present study (Foyer et al., 2009; Foyer and Noctor, 2009; Heber, 2002). Segarra et al. (2010) reported that one of the first responses of tomato plants in diseases caused by *Fusarium* is stomatal closure; with this reaction, the plant seeks to keep its water state stable under different conditions; however, if these conditions are prolonged, alterations occur in other processes. Our study suggests that in the treatments where the plants were inoculated with *Fol*, water limitations occurred due to vascular obstructions, so the plants responded by closing their stomata, avoiding water loss.

The increase in the concentration of proline in the tissues 15 DAI seems to be more an indicator of severe stress as reported in other studies (Alsamir et al., 2017; Mona et al., 2017), the authors use the proline contents as stress indicators, directly relating the severity of the symptoms with the content of proline. In agreement with the gs and RWC parameters, the increase in proline content occurred as a response to water shortage to maintain an osmotic adjustment in the leaves. In this context, the reductions in gs and RWC found in *Fol* infected tomato plants cause a decrease in energy flow observed in the fluorescence variables during the development of vascular wilt. In this way, the xylematic water potential decreases and induces stomatal closure; this, in turn, leads to a reduction in the photosynthetic rate due to limitations in CO_2_ and water intake (Nankishore and Farrell, 2016; Walters, 2015; Pshibytko et al., 2006).

Several components may be related to the decrease in chlorophylls during *Fol* infection. On the one hand, the oxidative cascades of the defense response, added to the accumulation of free radicals in the chloroplast after stomatal closure, can induce membrane damage by lipid peroxidation; these damages include the components of the photosynthetic apparatus such as thylakoid membranes and photosystems. Furthermore, the production of fungal toxins such as fusaric acid increases ROS production, and the degradation of membranes exacerbates oxidative damage and also damages to chloroplasts (Narula et al., 2020; Singh et al., 2017). All these components could contribute to damaging the photosystems of the plants infected with *Fol*, as observed in the results of the fluorescence parameters reported in this work.

The maintenance of chlorophylls integrity by chitosan suggests that after its application, protection mechanisms of the apparatus and photosynthetic pigments are activated in the plant. Similar results were obtained in other studies where the photosynthetic performance (30-60%) and the concentrations of chlorophylls and carotenoids were higher (30-74%) in plants that were treated with chitosan compared to their controls (Van et al., 2013; Dzung et al., 2011). According to our results, the chitosan application activates protective mechanisms such as the accumulation of proline, contributing in part to the protective capacity of membranes and proteins (Luan et al., 2018; Soliman end El-Mohamedy, 2017; Saharan and Pal, 2016). All the defense processes of the plant require the use of energy so that in addition to the direct alterations of the pathogen in the plant (i.e., water flow, CO_2_ uptake and assimilation, oxidative stress, and photosynthetic damage) (Yadeta and Thomma, 2013), the metabolic cost of defense due to the diversion of resources causes a significant decrease in biomass accumulation (Huot et al., 2014).

The results obtained in the expression of defense genes are congruent with those reported in other works where the expression of various defense genes occurs after 72 hours of treatment with chitosan and other resistance inducers in tomato plants (Jamiołkowska, 2020). These results were related to a lower expression of the disease caused by *R. solanacearum* and *Fol*, indicating that chitosan induces priming of plant defenses (Zehra et al., 2017; Kiirika et al., 2013).

The *PR1a* protein is induced during SA biosynthesis and is a SAR (Systemic Acquired Resistance) response marker (Jia et al., 2016; Iriti and Faoro, 2009). In other pathosystems such as tomato-*Fusarium andiyazi*, chitosan activates the *PR1* and *SOD* genes, indicating SA-mediated defense and antioxidant response co-occur (Chun and Chandrasekaran, 2019); similarly, in the kiwi-*Pseudomonas syringae* pv. *actinidiae* pathosystem, the application of chitosan induced 3.5 times the expression of the *PR1* gene when compared to the control (Corsi et al., 2017). The gene expression results indicate that chitosan presumably induces the plant to strengthen its immune system against pathogens, highlighting the potential of chitosan as a preventive, biocompatible, and non-toxic strategy for disease management (Dubin et al., 2020; Maluin and Hussein, 2020; Chowdhury et al., 2017).

The alterations in plant processes caused by *Fol* infection and its mitigation by chitosan are depicted in Fig. 9. During *Fol* infection, one of the first effects on the plant is the difficulty in taking water and stomatal closure, causing less assimilation of CO_2_ and damage to the photosystems. The plant increases its proline content due to water stress. In the presence of chitosan, plants activate the *PR1a, LOXA*, and *ERF1* genes upon perceiving infection; furthermore, plants also alleviate the hydric state, maintaining gs and CO_2_ assimilation and reducing oxidative damage; this may be due, in part, to proline, but also to other mechanisms not studied in this work that maintain the integrity of the photosynthetic system.

**Figure 9.**
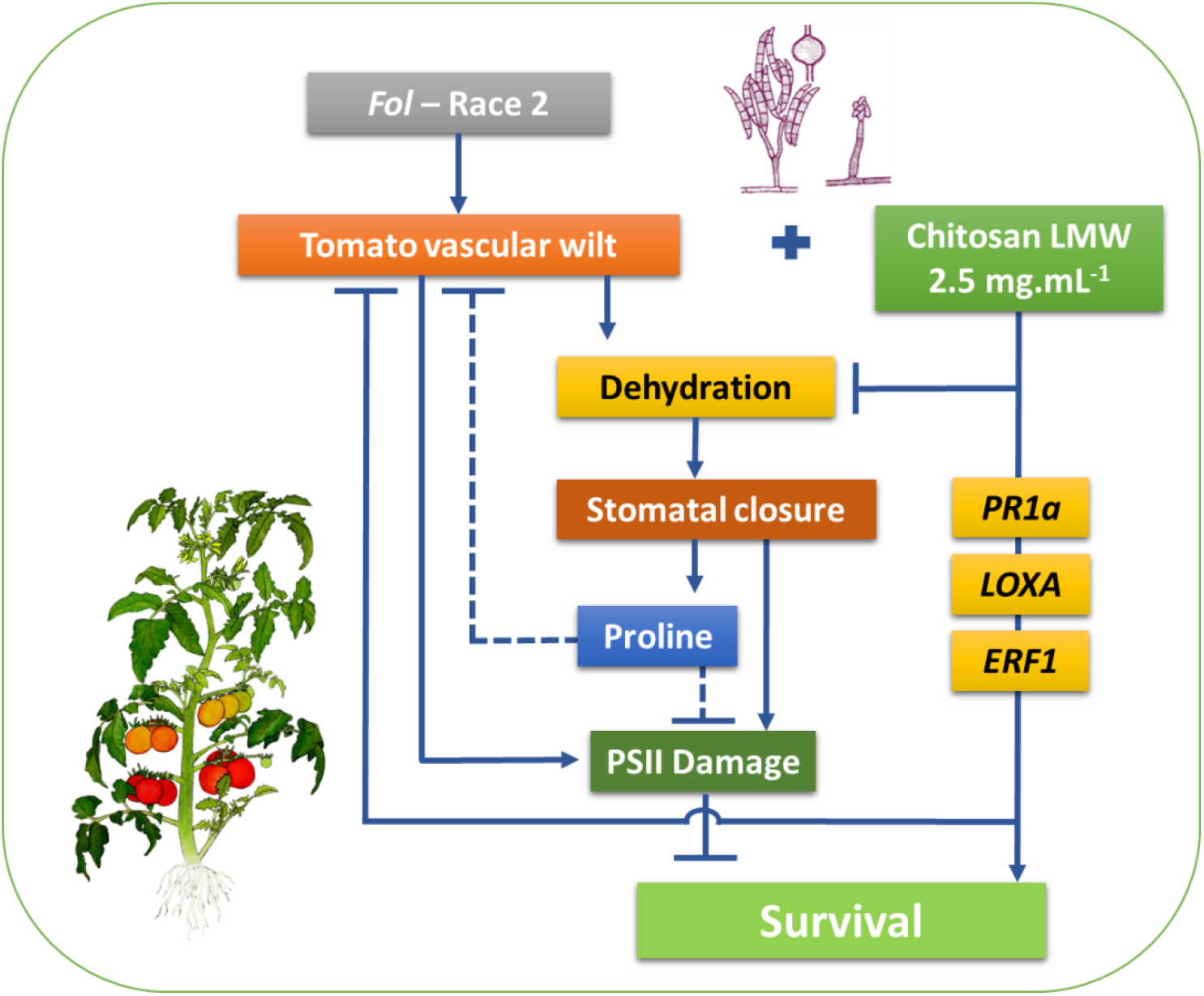
Model that describes the induction of resistance by chitosan in the tomato-*Fol* pathosystem. Truncated lines indicate interrupted processes; dashed lines refer to mechanisms hypothesized according to the results of the present study. Source: elaborated by the authors.

## Conclusions

The disease control capacity expressed by chitosan LMW (2.5 mg.mL^−1^), under favorable conditions for the establishment of the pathogen, reveals that this molecule has a high control potential against *Fol*. The physicochemical characteristics of chitosan and its biodegradability, together with the results presented here, making chitosan an excellent candidate for its use in sustainable disease management and clean agricultural production.

## Abbreviations

ABA: abcisic acid
AUDPC: area under the disease progress curve
JA: jasmonic acid
SA: salicylic acid
ATP: adenosine triphosphate
COM_2_: carbon dioxide
RWC: relative water content
ERF1: ethylene response factor1
ET: Ethylene
*Fol*: *F. oxysporum* f. sp. *lycopersici*
*Forl*: *F. oxysporum* f. sp. *radicis lycopersici*
Fv/Fm: maximum potential quantum efficiency of PSII photochemistry
gs: stomatal conductance
HMW: high molecular weight
H_2_O_2_: hydrogen peroxide
LMW: low molecular weight
LOXA: lipoxygenase A
MAMPs or PAMPs: Microbial or Pathogen Associated Molecular Patterns
MMW: medium molecular weight
MAPKs: Mitogen-activated protein kinase
NADPH^+^: nicotinamide adenine dinucleotide phosphate
NO: nitric oxide
PAL: phenylalanine ammonia lyase
PDA: potato dextrose agar
PIRG: percentage inhibition of radial growth
PR: Pathogenesis-related
PR1a: pathogen response1
PSI: Photosystem I
PSII: Photosystem II
PTI: PAMP Triggered Immunity
Q_A_: quinone A
qP: photochemical quenching
qRT-PCR: quantitative real time polymerase chain reaction
ROS: reactive oxygen species
RuBisCO: ribulose-1,5-bisphosphate carboxylase/oxygenase
Y(II): photochemical efficiency of PSII
ΔCT: delta cycle threshold

## Author’s contributions

Conceptualization: MS-S, SLC and ETR. Methodology: SLC, AVN, DBD, and MGM. Formal analysis: SLC, AVN, DBD, MGM, ETR, and MS-S. Writing-original draft preparation: SLC, AVN, DBD, MGM, ETR, and MS-S. Project administration: MS-S. All authors read and approved the final manuscript.

## Acknowledgments

This work was supported by the Corporación Colombiana de Investigación Agropecuaria, AGROSAVIA, and the Ministerio de Agricultura y Desarrollo Rural (MADR).

## Data availability statement

The data that supports the findings of this study are available in the supplementary material of this article. Any additional data will be available on request to the corresponding author.

## Notes

### Competing Interest Statement

The authors have declared no competing interest.

